# The Role of *fosA* in Challenges with Fosfomycin Susceptibility Testing of Multispecies *Klebsiella pneumoniae* Carbapenemase-Producing Clinical Isolates

**DOI:** 10.1101/611970

**Authors:** Zachary S. Elliott, Katie E. Barry, Heather L. Cox, Nicole Stoesser, Joanne Carroll, Kasi Vegesana, Shireen Kotay, Anna E. Sheppard, Alex Wailan, Derrick W. Crook, Hardik Parikh, Amy J. Mathers

**Affiliations:** Department of Pharmacy Services, University of Virginia Health System, Charlottesville, Virginia, USA.; Division of Infectious Diseases and International Health, Department of Medicine, University of Virginia Health System, Charlottesville, Virginia, USA.; Modernizing Medical Microbiology Consortium, Nuffield Department of Clinical Medicine, John Radcliffe Hospital, Oxford University, Oxford, United Kingdom.; NIHR Health Protection Research Unit in Healthcare Associated Infection and Antimicrobial Resistance at University of Oxford in partnership with Public Health England, Oxford, UK.; Clinical Microbiology, Department of Pathology, University of Virginia Health System, Charlottesville, Virginia, USA.; Health Information & Technology, University of Virginia Health System, Charlottesville, Virginia, USA; School of Medicine Research Computing, University of Virginia, Charlottesville, USA

**Author notes:** **Corresponding Author:** Amy J. Mathers, MD, D(ABMM), P.O. Box 800255, Charlottesville, VA, USA 22908-1361.

## Abstract

With multidrug resistant (MDR) Enterobacteriales on the rise, a non-toxic agent with a unique mechanism of action such as fosfomycin seems attractive. However, establishing accurate fosfomycin susceptibility testing for non-*E. coli* in a clinical microbiology laboratory remains problematic. We evaluated fosfomycin susceptibility by multiple methods with multiple strains and species of KPC-producing clinical isolates collected at a single center between 2008 and 2016. In addition, we assessed the presence of fosfomycin resistance genes from whole genome sequencing (WGS) data using NCBI’s AMRFinder and custom HMM search. Susceptibility testing was performed using glucose-6-phosphate supplemented fosfomycin E-Test and Kirby-Bauer disk diffusion (DD) assays, and compared to agar dilution. Clinical Laboratory and Standards Institute (CLSI) breakpoints for *E. coli* were applied for interpretation. Overall, 63% (60/96) of isolates were susceptible by E-Test, 70% (67/96) by DD, and 88% (84/96) by agar dilution. *FosA* was detected in 80% (70/88) of previously sequenced isolates, with species-specific associations and alleles, and *fosA*-positive isolates were associated with higher MIC distributions. Disk potentiation testing was performed using sodium phosphonoformate to inhibit *fosA* and showed significant increases in the zone diameter of DD testing for isolates that were *fosA*-positive compared to *fosA*-negative. The addition of sodium phosphonoformate (PPF) corrected 10/14 (71%) major errors in categorical agreement with agar dilution. Our results indicate that *fosA* influences the inaccuracy of susceptibility testing by methods readily available in a clinical laboratory when compared to agar dilution. Further research is needed to determine the impact of *fosA* on clinical outcomes.

## Introduction

Antimicrobial resistance among gram-negative organisms continues to increase and presents a serious threat to modern medicine with carbapenemase-producing *Enterobacteriaceae* (CPE) considered one of the most pressing issues (1). The concern with CPE is largely due to a lack of remaining therapeutic options, especially oral agents (2). This has led to the re-evaluation of older antimicrobials to combat infections due to multidrug-resistant (MDR) Gram-negative pathogens. Fosfomycin, which was originally discovered in 1969, has been shown to have *in vitro* activity against CPE (3). In the United States, the oral formulation is available for the treatment of uncomplicated urinary tract infections due to susceptible strains of *Escherichia coli* and *Enterococcus feacalis.* Outside of the United States, the intravenous (IV) formulation is approved and available for the management of systemic infections (4). Zavante Pharmaceuticals received Fast Track designation from the FDA in 2015 for IV fosfomycin and have completed Phase III clinical trials for the United States market (5).

Fosfomycin is a bactericidal antibiotic that binds to the cysteine residue of UDP-*N*-acetylglucosamine enolpyruvyl transferase (MurA) and inhibits peptidoglycan biosynthesis (6). Fosfomycin has activity against a range of bacterial pathogens, including highly drug resistant *Enterobacteriaceae* (6). Fosfomycin resistance in Enterobacteriales has been primarily driven by mutations in the *glpT* and *uhpT* genes, preventing active transport of fosfomycin into the cell (7). These mutations are however thought to be associated with a fitness cost in *E. coli* and are thus unstable (8, 9). The other major mechanism of resistance is hydrolysis of the drug via diverse Fos enzymes: FosA (FosA2, FosA3, FosA4, FosA5, FosA6, FosA7), FosB, and FosX are metalloenzymes, whereas FosC is a serine enzyme (10). FosA was originally discovered on a transposon, Tn2921, in a *Serratia marcescens* plasmid, and catalyzes the addition of glutathione to fosfomycin, rendering the drug ineffective (11). Transmissible *fosA* is of most concern in Enterobacteriales, and plasmid-mediated *fosA3* has been increasingly identified in *E. coli* in Europe (12). A recent evaluation of Fos enzymes in non-*E. coli* Enterobacteriales demonstrated different *fosA* variants were shown to be chromosomally located in a species-specific manner (11) Lastly, MurA target site alternation can also confer fosfomycin resistance. Amino acid substitutions in MurA, most notably Asp369Asn and Leu370lle, have been responsible for fosfomycin resistance (13).

Susceptibility testing of fosfomycin for non-*E. coli* Enterobacteriales is difficult for clinical microbiology labs (14–16). Both CLSI and EUCAST specifically recommend against the use of broth microdilution methods which likely impacts the inaccuracies with most automated susceptibility testing platforms for *E. coli* or *Klebsiella pneumoniae* (17, 18). Agar dilution is considered the reference method, and endorsed by EUCAST, however this is difficult to execute routinely in a clinical microbiology laboratory. Kirby-Bauer Disk Diffusion (DD) and E-Tests are more attractive options, as they can be performed easily in a clinical laboratory, but colonies often grow within the zones of inhibition making interpretation difficult (19). Attempting to change the zone cutoff to better align with agar dilution has not proved successful in non-*E. coli* Enterobacteriales (15). With agar dilution as the only accurate method for non-*E. coli* Enterobacteriales, we aimed to characterize some of the molecular mechanisms (by whole genome sequencing for a subset of isolates) that may be contributing to the inaccuracies with these diffusion methods, using a set of clinical, non-*E. coli* carbapenemase producing strains and agar dilution-based reference phenotyping.

## Materials and Methods

Retrospective samples of *Klebsiella pneumoniae* carbapenemase KPC-producing *Gammaproteobacteria* isolates were selected from those collected at University of Virginia Health System since August 2008. Isolates were chosen to represent diverse species and strains for which Illumina sequence data were available. Species identification had been performed by matrix-assisted laser desorption ionization-time of flight mass spectrometry (MALDI-TOF, VITEK-MS, VITEK-2, bioMêrieux) and all isolates were *bla*_KPC_ PCR positive as previously described (20).

E-Test was performed using glucose-6-phosphate (G6P) supplemented fosfomycin E-Test strips (BioMêrieux Durham, NC) according to the manufacturer instructions. DD was performed with a 200 µg fosfomycin disk with 50µg G6P (Becton and Dickinson, Franklin Lakes, NJ) on Mueller-Hinton agar as per CLSI guidelines(21). Agar dilution was performed using G6P supplemented (25 µg/mL) Mueller-Hinton agar with fosfomycin concentrations ranging from 0.5 to 1,024 µg/mL per CLSI methods(22). A 0.5 McFarland inoculum for each isolate was placed in triplicate on the agar, placed in an incubator at 35^0^ C for 16 to 20 hours, and then interpreted. Plates were prepared without isolates at each concentration to serve as the control.

Per the package insert for interpretation of E-Test, the crossing point of the ellipse was used to identify the minimum inhibitory concentration (MIC) where colonies within the zone of the ellipse were accounted for if 5 colonies were present within 3mm of the strip within the zone (BioMêrieux Durham, NC). DD diameters were measured as the shortest distance between 2 separate colonies (17, 21). For agar dilution, the median interpreted MIC was recorded as the result. All fosfomycin susceptibilities were interpreted according to CLSI breakpoints for *E. coli* urinary isolates as there are no breakpoints available for non-*E. coli* Enterobacteriales (17).

With agar dilution as the reference method, essential agreement for E-test and DD were defined as MIC variation within on dilution. Categorical agreement was defined as matching susceptible/intermediate/resistant interpretation criteria for the two respective tests, as per CLSI guidelines for *E. coli* urinary isolates. Falsely susceptible results were deemed to be very major errors and falsely resistant results to be major errors. All other disagreements were deemed minor errors. Chi-Square and Fisher’s Exact Tests were used to compare rates of non-susceptibility and categorical agreement. The Mann-Whitney test was utilized for statistical analysis of MIC distributions and DD zone diameter changes.

Disk potentiation testing with sodium phosphonofomate (PPF) was performed to specifically evaluate the activity of *fosA* enzymes and the impact on susceptibility testing with the disk diffusion method. Cultures of each isolate were plated on Mueller-Hinton agar with 1mL of a 50mg/mL sodium phosphonoformate (PPF; Sigma-Aldrich) solution to a 200 µg fosfomycin disk supplemented with 50µg G6P. The plates were incubated overnight at 37^0^ C and the inhibition zone was recorded. These inhibition zones were then compared the DD inhibition zones of fosfomycin without the supplementation of PPF. For each isolate tested, a blank disk with PPF was also placed on the agar plate to serve as a negative control.

Molecular mechanisms of resistance to fosfomycin were investigated in a subset of isolates previously whole genome sequenced by Ilumina Sequencing (HiSeq, 2000) as previously described (20). The quality filtered short reads were *de novo* assembled using SPAdes v3.11 (23), and the contigs were screened for *fos* resistance genes using NCBI’s AMRFiner (id ≥ 0.9; cov ≥ 0.5)(24). For isolates where AMRFinder failed to detect *fos* genes, we screened the contigs using a custom HMM model built from distinct *fosA* protein sequences published by Ito *et. al.*, with an e-value threshold of 1e-20 (11, 25).

## Results

### Fosfomycin Susceptibility across Species

Ninety-six *bla*_KPC_-positive isolates across twelve species were included in the study (Table 1 for species breakdown). Eighty-eight of the 96 isolates had undergone whole genome sequencing (WGS). The MIC_50_ across all isolates was 8 µg/mL and the MIC_90_ was 128 µg/mL by agar dilution. Using the 2019 CLSI breakpoints (≤64 µg/mL=S), 84 of 96 isolates (88%) were susceptible, 11 of 96 (11%) were resistant, and 1 of 96 (1%) was classified as intermediate. The MIC distributions by agar dilution are shown in Figure 1.

**Table 1:**
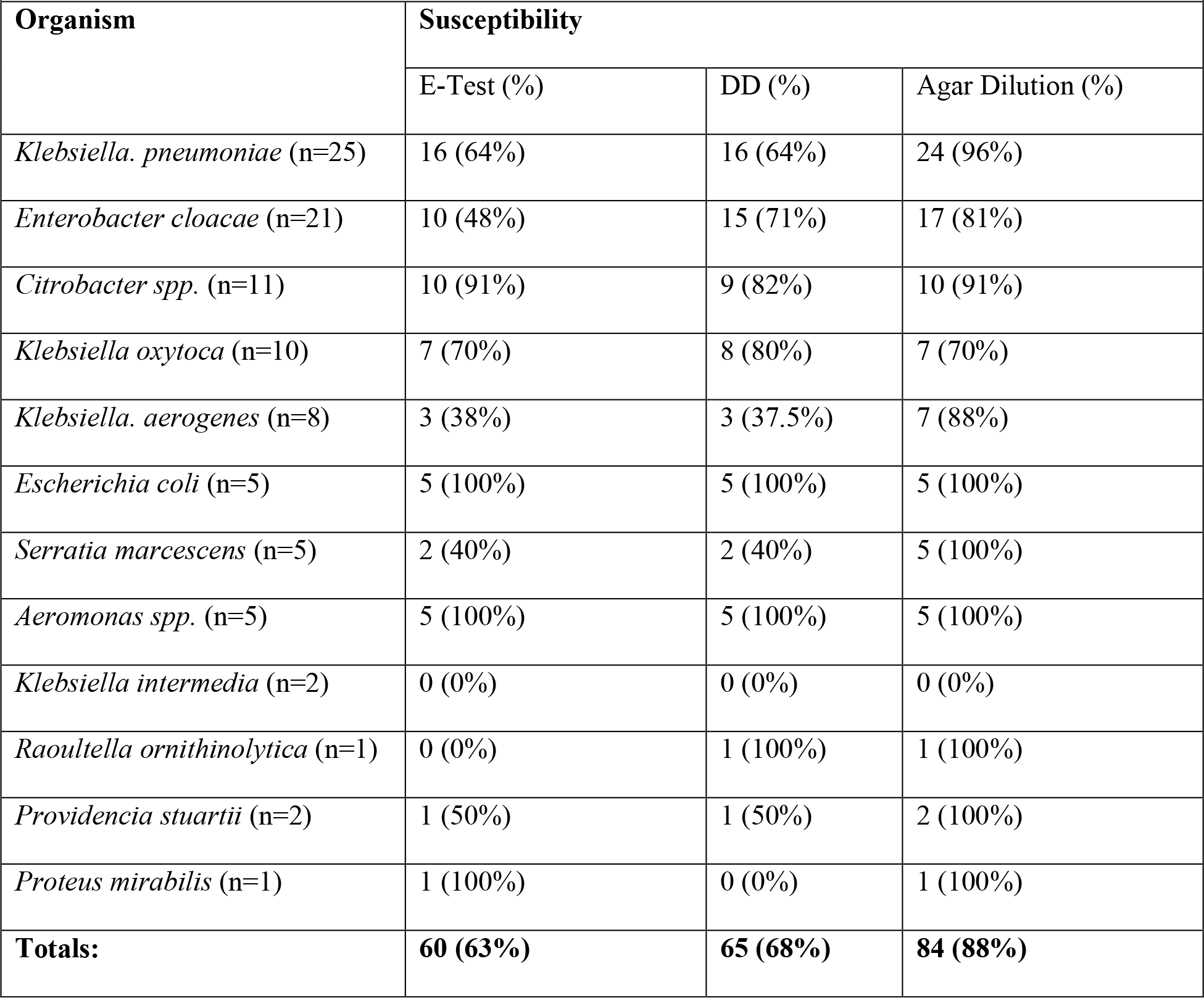
Fosfomycin Susceptibility (n=96)

Appendixes
| **Table 1: Fosfomycin Susceptibility (n=96)** |  |  |  |
| --- | --- | --- | --- |
| **Organism** | **Susceptibility** |  |  |
|  | E-Test (%) | DD (%) | Agar Dilution (%) |
| *Klebsiella. pneumoniae* (n=25) | 16 (64%) | 16 (64%) | 24 (96%) |
| *Enterobacter cloacae* (n=21) | 10 (48%) | 15 (71%) | 17 (81%) |
| *Citrobacter spp.* (n=11) | 10 (91%) | 9 (82%) | 10 (91%) |
| *Klebsiella oxytoca* (n=10) | 7 (70%) | 8 (80%) | 7 (70%) |
| *Klebsiella. aerogenes* (n=8) | 3 (38%) | 3 (37.5%) | 7 (88%) |
| *Escherichia coli* (n=5) | 5 (100%) | 5 (100%) | 5 (100%) |
| *Serratia marcescens* (n=5) | 2 (40%) | 2 (40%) | 5 (100%) |
| *Aeromonas spp.* (n=5) | 5 (100%) | 5 (100%) | 5 (100%) |
| *Klebsiella intermedia* (n=2) | 0 (0%) | 0 (0%) | 0 (0%) |
| *Raoultella ornithinolytica* (n=1) | 0 (0%) | 1 (100%) | 1 (100%) |
| *Providencia stuartii* (n=2) | 1 (50%) | 1 (50%) | 2 (100%) |
| *Proteus mirabilis* (n=1) | 1 (100%) | 0 (0%) | 1 (100%) |
| **Totals:** | **60 (63%)** | **65 (68%)** | **84 (88%)** |

**Table 2.** MIC Distributions Among Most Frequent Isolates

| Table 2. MIC Distributions Among Most Frequent Isolates |  |  |  |  |  |  |
| --- | --- | --- | --- | --- | --- | --- |
| Species (n) | FosA<br>Positive* | MIC data (mcg/mL) |  |  | Nonsusceptibility |  |
|  |  | Range (mcg/mL) | MIC<sub>50</sub> | MIC<sub>90</sub> | No. of Isolates | % |
| All species (96) | 70/88<br>(79.55%) | ≤ 0.5 to >1024 | 16 | 128 | 12 | 12.5% |
| *K. pneumoniae* (25) | 24/24<br>(100.00%) | 2 to 512 | 8 | 32 | 1 | 4.0% |
| *E. cloacae* (21) | 19/21<br>(90.48%) | ≤ 0.5 to >1024 | 16 | 512 | 4 | 19.05% |
| *Citrobacter freundii* (10) | 2/10 (20.00%) | ≤ 0.5 to >1024 | 0.5 | 16 | 1 | 10.0% |

\* Isolates with WGS data available

**Figure 1.**
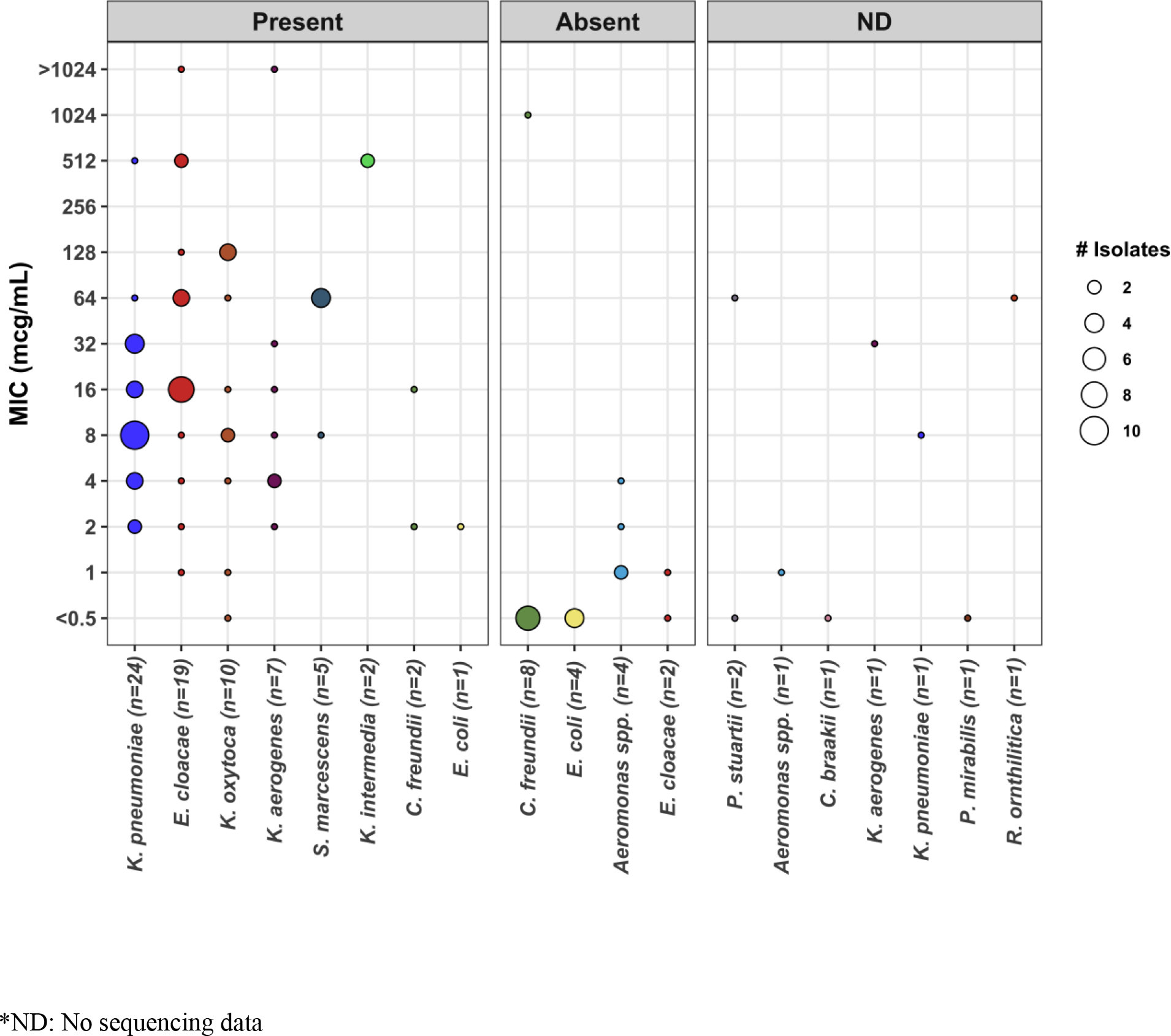
MIC distribution of KPC-producing isolates, grouped by *fosA* resistance gene presence screened from whole-genome sequencing data.

### Diffusion Method Performance

Sixty of 96 isolates (63%) were susceptible by E-Test and 65/96 (68%) were susceptible by DD. Categorical agreement of E-Test to agar dilution occurred in 69/96 isolates (72%) with 1 very major error, 16 major errors, and 10 minor errors. Essential agreement occurred in 55 of 96 isolates (57%) overall and in 4 of 5 (80%) *E. coli* isolates. Categorical agreement of DD to agar dilution occurred in 72/96 isolates (75%) with 2 very major errors, 14 major errors, and 8 minor errors. Of note, when testing the non-*E. coli* species, colonies within the zone were frequently present thus making interpretation challenging but adhered to package insert and CLSI guidance for E-test and DD respectively(21).

### FosA presence

Of the isolates with WGS data, no isolate harbored *fosC*, while 70/88 isolates (80%) harbored an allele of *fosA*. All *K. pneumoniae* isolates (*n*=24) carried *fosA*, with 23 of 24 isolates (96%) harboring the *fosA6* or *fosA6-*like variant. Interestingly, only *K. aerogenes* isolates (n=7; 100%) carried the same variant. Detection of *fosA* in all tested isolates is shown in Figure 1. Isolates without *fosA* (*n*=18) had a MIC range of ≤0.5 to 1024 µg/mL, MIC_50_ of ≤0.5 µg/mL, and MIC_90_ of 2 µg/mL. One of the 18 isolates (5.56%) was non-suseptible to fosfomycin. Isolates harboring *fosA* (*n*=70) had a MIC range of ≤0.5 to >1024 µg/mL, MIC_50_ of 16 µg/mL, and MIC_90_ of 128 µg/mL. Eleven of the 70 isolates (16%) were non-susceptible to fosfomycin. Isolates carrying the *fosA* gene were associated with a higher MIC distribution as compared to those without the gene (*P*=<0.00001), but did not differ in rates of non-susceptibility (*P*=0.26). These results are shown in Table 3.

**Table 3.** Comparison of Isolates

| **Table 3. Comparison of Isolates** |  |  |  |  |  |  |
| --- | --- | --- | --- | --- | --- | --- |
| Isolates | Range | MIC<br>50 | MIC<br>90 | *P*-value | No. of nonsusceptible<br>isolates (%) | *P*-value |
| Not harboring *fosA*<br>(n=18) | ≤0.5 to<br>1024 | ≤0.5 | 2 | ***P* < 0.00001** | 1 (5.56%) | ***P* = 0.2627** |
| Harboring *fosA* (n=70) | ≤0.5 to<br>>1024 | 16 | 128 |  | 11 (15.71%) |  |

### FosA inhibition effect on Susceptibility Testing by Diffusion Method

Disk potentiation testing with PPF was performed on all 96 isolates. Categorical agreement with agar dilution was found in 72/96 (75%) of isolates prior to the additional of PPF and in 81/96 (84%) isolates after the addition of PPF (p=0.11). In the 88 isolates with WGS available, rates of categorical agreement were compared with and without PPF. In isolates that were negative for the *fosA* gene, categorical agreement was found in 18/18 (100%) and 17/18 (94%) of isolates before and after the additional of PPF, respectively (p=1). In isolates that carried the *fosA* gene, categorical agreement was found in 49/70 (70%) and 59/70 (84%) before and after the addition of PPF, respectively (p=0.04). When specifically isolating all non-susceptible isolates by DD, categorical agreement was found in 9/31 (29%) isolates and 20/31 (65%) before and after the addition of PPF, respectively (p=0.005). Results are shown in Table 4 and Table 5. The presence of PPF not only increased the zone size but also greatly decreased the presence of colonies within the zone in DD testing for the *fosA-*positive isolates (Figure 2).

**Table 4.** Disk Potentiation testing on all isolates (n=96)

| **Table 4. Disk Potentiation testing on all isolates (n=96)** |  |  |  |  |  |
| --- | --- | --- | --- | --- | --- |
| DD Fosfomycin Only |  |  | DD Fosfomycin + PPF |  |  |
| Susceptibility | n= | Categorical agreement | Susceptibility | n= | Categorical agreement |
| Susceptible | 65 | 63/65=96.9% | Susceptible | 80 | 76/80=95.0% |
| Intermediate | 7 | 0/7=0% | Intermediate | 6 | 0/6=0% |
| Resistant | 24 | 9/24=37.5% | Resistant | 10 | 5/10=50% |
|  |  | **Total: 72/96=75%** |  |  | **Total: 81/96=84%** |
| ***P* = 0.10644** |  |  |  |  |  |

**Table 5.**
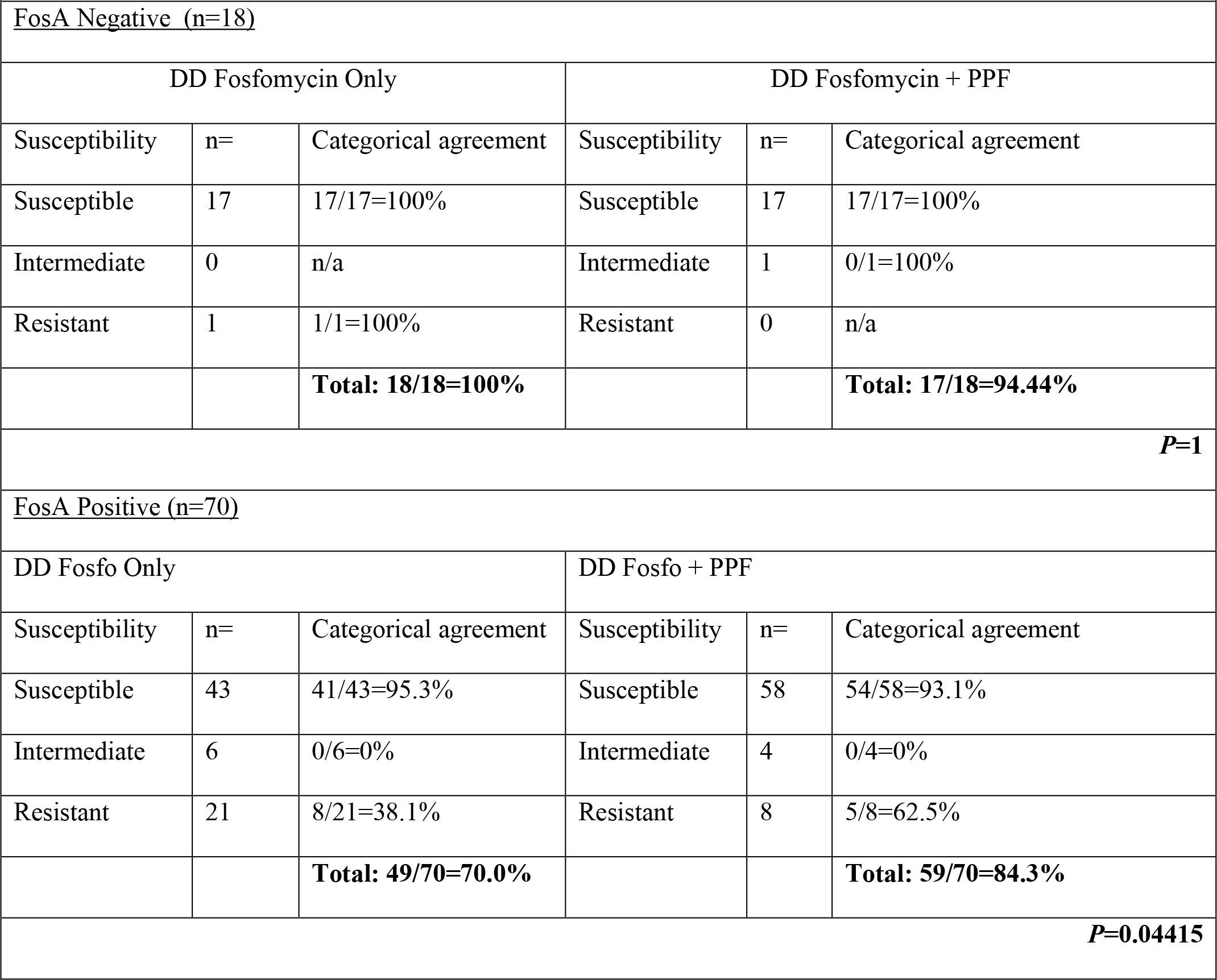
Disk Potentiation testing on WGS Isolates (n=88)

**Figure 2.**
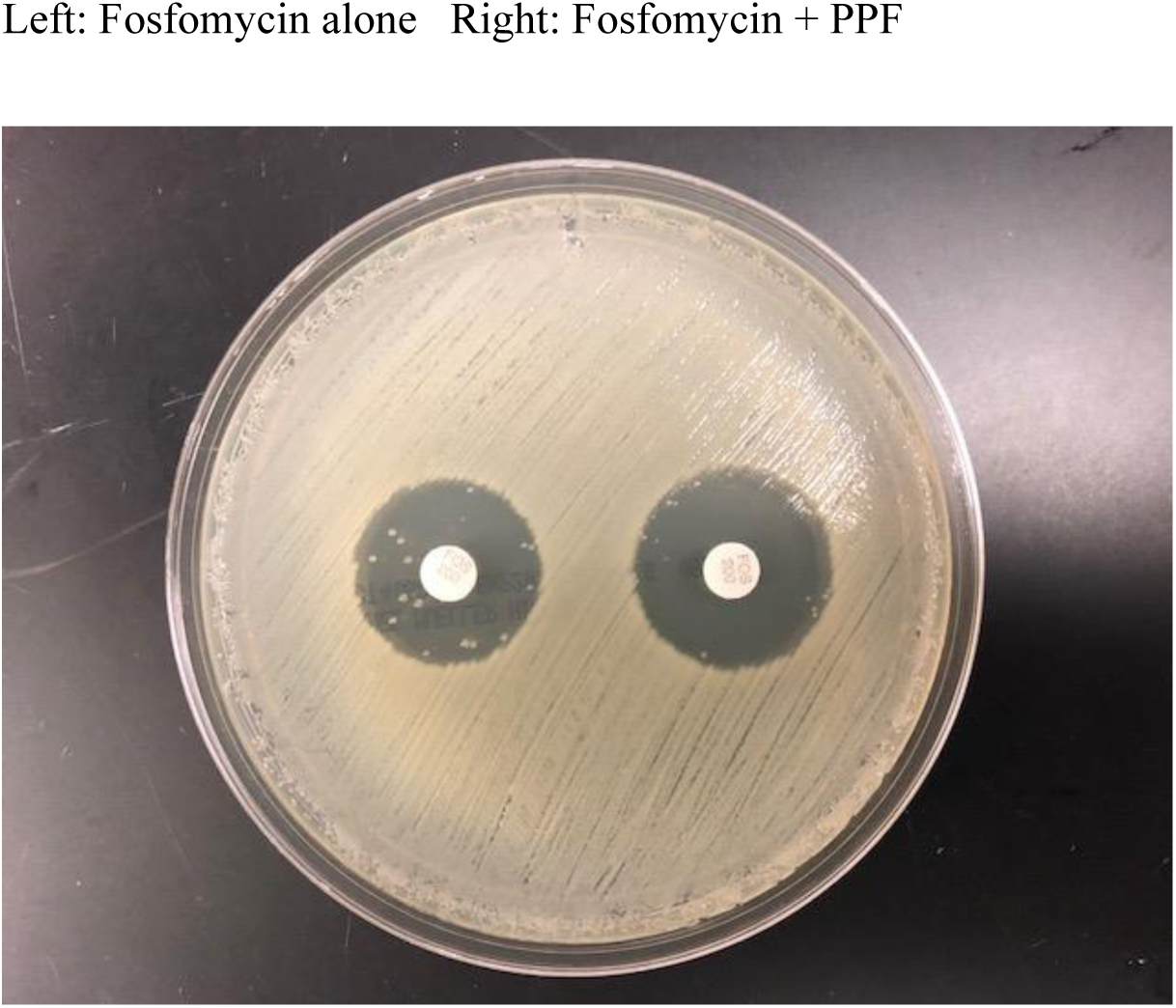
Example of elimination of fosfomycin nonsusceptible sub-colonies within zone of inhibition in *Klebsiella pneumonia* CAV 1217

## Discussion

We demonstrate that fosfomycin susceptibility testing by routinely used laboratory diffusion-based methods (E-test and DD) largely overcalls resistance, when compared to agar dilution as a gold standard (see Table 1). Fosfomycin susceptibility appears to be influenced by the presence of *fosA* among these KPC-producing Enterobacteriales isolates. This is highly relevant to the clinical microbiologist who is frequently fielding requests for fosfomycin susceptibility testing for non-*E. coli* Enterobacteriales. A prior study by Kaase *et al.* tested 107 carbapenem-non-susceptible *Enterobacteriaceae* isolates, of which 80 produced various carbapenemases (KPC, VIM, NDM, OXA-48) and similarly found 81% of isolates to be susceptible to fosfomycin with a MIC of ≤64 mcg/mL by agar dilution. This study also found similar issues of discordance with diffusion testing methods (15), as was also seen in Hirsch *et al*. (14).

Fosfomycin has been promoted as a useful, safe medication for the treatment of urinary tract infections against multidrug resistant non-*E. coli Enterobacteriaceae* (2, 3, 7, 26, 27) but susceptibility testing by agar dilution is practically difficult for a clinical microbiology lab. The CLSI cutoffs for *E. coli* were applied for E-test and DD interpretation for all species tested in this experiment, which requires accounting for scattered colonies within the ellipse per the package insert or zone per CLSI (17). Fosfomycin susceptibility testing for *E. coli* was recently reviewed by CLSI in 2018. At that time the recommendation that susceptibility cut-offs only apply to *E. coli* was strengthened and, since scattered colonies are rare within the zone for this species the practice of measuring the zone from the innermost colonies was upheld (17). This differs from the new European Committee on Antimicrobial Susceptibility Testing (EUCAST) guidelines for fosfomycin and *E. coli*, which suggests ignoring scattered colonies within diffusion-based inhibition zones, and utilizing agar dilution approaches for non-*E.coli* Enterobacteriales (28). This decision for the former was based on the findings that sub-colonies of *E. coli* within inhibition zones are rare (<1% of isolates) and largely less fit with channel or transporter mutations (8, 9).

We postulate based on our findings that the colonies within the inhibition zone seen more frequently with some non-*E. coli* species may be driven by the presence of a chromosomal *fosA* rather than channel or transporter mutations. Thus the advice to ignore the colonies in the zone in *E. coli* may not apply to non-*E. coli* Enterobacteriales and clinical microbiologists should proceed with caution. The discordance among commercially available DD and E-test with agar dilution we observed was largely due to bacterial colonies that grow within the zone of inhibition. Based on the change in zone size with the addition of a FosA inhibitor as well as the work of others demonstrating the activity of chromosomal expressed FosA, it may unwise to ignore sub-colonies as the clinical implications of this finding remain unknown (29).

In our subset of *E. coli* isolates (n=5), categorical agreement was found in 5 of 5 (100%) isolates. All *E. coli* isolates were susceptible to fosfomycin by both DD and agar dilution, and no colonies were observed within the zones of inhibition. Although our numbers are small, this is consistent with other reports (8).

*FosA* (alleles 1-7) was identified in the majority of the clinical isolates in this study. *FosA* was present in all *Klebsiella spp.* isolates, with a large portion harboring the *fosA6* allele*. FosA6* was first reported in 2016, from an ESBL-producing, fosfomycin-resistant *E. coli* strain in Pennsylvania, USA. It shared 96% identity with *fosA5* and 79% identity with *fosA3* but was located on a plasmid, unlike the chromosomally-encoded *fosA* in *K. pneumoniae* (30). It has been suggested that *fosA6* was mobilized from the chromosome of *K. pneumoniae* to an *E. coli* plasmid (30). However, in our study, no *E. coli* harbored *fosA6*, but rather one isolate harbored *FosA7*, which has been described on the chromosome of *Salmonella enterica* (31). We postulate that this gene may have been acquired via plasmid transfer with *Salmonella enterica* serving as the reservoir for this allele. No *fosC* was detected, as expected, as it is a gene found most commonly within *Pseudomonas* spp. which were not included in this study.

In this subset of isolates, the presence of *fosA* resulted in a trend towards higher MIC values compared to isolates not harboring the gene. Despite the higher distribution of MIC values, there was no statistical difference in fosfomycin susceptibility when performed by the agar dilution method within the range of CLSI range of susceptible for *E. coli*.

Disk potentiation testing with PPF was performed to specifically evaluate the activity of *fosA* enzymes and the impact on susceptibility testing with DD. The addition of PPF significantly increased zone diameter size and, subsequently, improved categorical agreement of disk diffusion with agar dilution, particularly in *fosA-*positive isolates in which most major errors were eliminated. These improvements were largely due to the elimination of sub-colonies within the zone of inhibition, as illustrated for isolate CAV 1217 (Figure 2). As expected, there was no statistical change in categorical agreement in isolates that did not harbor a *fosA* allele. The results of the disk potentiation testing indicate that *fosA* impacts fosfomycin activity and limits rapid, diffusion-based susceptibility testing. Some isolates had substantial zone diameter increases (6-8mm) after the addition of PPF, yet did not alter the susceptibility interpretation. This is likely due to alternative mechanisms of fosfomycin resistance, such as transporter mutations or MurA mutations. An alternative explanation is that certain alleles of *fosA* may have the ability to overcome the inhibition of PPF that was added to the disk.

Lastly, our data suggests that the addition of PPF may have a synergistic effect with fosfomycin against *fosA-*positive organisms. This is corroborated by a recently published study that found significant MIC reductions and restored fosfomycin susceptibility in *fosA-*positive Gram-negative organisms (29). PPF is available as the antiviral foscarnet but would likely be unattractive as an adjuvant therapy due to toxicity.

Our study is limited by a small sample size of *bla*_KPC_-positive multidrug-resistant isolates collected from a single center. However, the clinical utility of fosfomycin is primarily against MDR isolates and our study highlights the difficulty in accurately providing fosfomycin AST for these organisms. A further limitation is the lack of exploration of other molecular mechanisms of resistance which were not evaluated in all isolates. Our study also lacks outcome data, and therefore we can make no conclusions on the clinical implications of fosfomycin susceptibility testing results.

In conclusion, fosfomycin appears to have reliable *in vitro* activity against KPC-producing Gram-negative organisms by agar dilution. However, methods readily available in a clinical microbiology laboratory, E-test and DD, generate frequent major errors for the same isolates, with the presence of *fosA* impacting the interpretation of these diffusion-based methods. Caution is advised when interpreting and releasing AST results derived from diffusion-based methods for non-*E. coli Enterobacteriales*. Regardless, further research is needed to establish correlations between antimicrobial susceptibility testing, *fosA* presence and clinical outcomes.

## Acknowledgements

We thank UVaMC Clinical Microbiology staff for collection of study isolates.

## References

1. Centers for Disease Control and Prevention (CDC). 2013. Antibiotic resistance threats in the United States. (CDC) CfDCaP, Atlanta, GA USA.

2. Morrill HJ, Pogue JM, Kaye KS, LaPlante KL. 2015. Treatment Options for Carbapenem-Resistant Enterobacteriaceae Infections. Open Forum Infect Dis 2:ofv050.

3. Endimiani A, Patel G, Hujer KM, Swaminathan M, Perez F, Rice LB, Jacobs MR, Bonomo RA. 2010. In vitro activity of fosfomycin against blaKPC-containing Klebsiella pneumoniae isolates, including those nonsusceptible to tigecycline and/or colistin. Antimicrob Agents Chemother 54:526–9.

4. Grabein B, Graninger W, Rodriguez Bano J, Dinh A, Liesenfeld DB. 2017. Intravenous fosfomycin-back to the future. Systematic review and meta-analysis of the clinical literature. Clin Microbiol Infect 23:363–372.

5. Kaye KS, Rice LB, Dane A, Stus V, Sagan O, Fedosiuk E, das A, Skarinsky D, Eckburg P, Ellis-Grosse EJ. 2017. Intravenous Fosfomycin (ZTI-01) for the treatment of Complicated Urinary Tract Infections (cUTI) Including Acute Pyelonephritis (AP): Results from a Multi-Center, Randomized, Double-Blind Phase 2/3 Study in Hospitalized Adults (ZEUS). Open Forum Infectious Diseases 4:S528.

6. Falagas ME, Vouloumanou EK, Samonis G, Vardakas KZ. 2016. Fosfomycin. Clin Microbiol Rev 29:321–47.

7. Michalopoulos AS, Livaditis IG, Gougoutas V. 2011. The revival of fosfomycin. Int J Infect Dis 15:e732–9.

8. Lucas AE, Ito R, Mustapha MM, McElheny CL, Mettus RT, Bowler SL, Kantz SF, Pacey MP, Pasculle AW, Cooper VS, Doi Y. 2018. Frequency and Mechanisms of Spontaneous Fosfomycin Nonsusceptibility Observed upon Disk Diffusion Testing of Escherichia coli. J Clin Microbiol 56.

9. Marchese A, Gualco L, Debbia EA, Schito GC, Schito AM. 2003. In vitro activity of fosfomycin against gram-negative urinary pathogens and the biological cost of fosfomycin resistance. Int J Antimicrob Agents 22 Suppl 2:53–9.

10. Silver LL. 2017. Fosfomycin: Mechanism and Resistance. Cold Spring Harb Perspect Med 7.

11. Ito R, Mustapha MM, Tomich AD, Callaghan JD, McElheny CL, Mettus RT, Shanks RMQ, Sluis-Cremer N, Doi Y. 2017. Widespread Fosfomycin Resistance in Gram-Negative Bacteria Attributable to the Chromosomal. MBio 8.

12. Mendes A, Rodrigues C, Pires J, Amorim J, Ramos Mh, Novais A, Peixe L. 2016. Importation of Fosfomycin Resistance fosA3 Gene to Europe., vol 22, p 346–348. Emerging Infectious Diseases

13. Takahata S, Ida T, Hiraishi T, Sakakibara S, Maebashi K, Terada S, Muratani T, Matsumoto T, Nakahama C, Tomono K. 2010. Molecular mechanisms of fosfomycin resistance in clinical isolates of Escherichia coli. Int J Antimicrob Agents 35:333–7.

14. Hirsch EB, Raux BR, Zucchi PC, Kim Y, McCoy C, Kirby JE, Wright SB, Eliopoulos GM. 2015. Activity of fosfomycin and comparison of several susceptibility testing methods against contemporary urine isolates. Int J Antimicrob Agents 46:642–7.

15. Kaase M, Szabados F, Anders A, Gatermann SG. 2014. Fosfomycin susceptibility in carbapenem-resistant Enterobacteriaceae from Germany. J Clin Microbiol 52:1893–7.

16. Perdigao-Neto LV, Oliveira MS, Rizek CF, Carrilho CM, Costa SF, Levin AS. 2014. Susceptibility of multiresistant gram-negative bacteria to fosfomycin and performance of different susceptibility testing methods. Antimicrob Agents Chemother 58:1763–7.

17. Clinical Laboratory and Standards Institute. 2019. m100-S29 Performance Standards for Antimicrobial Susceptibility Testing, 29th ed. CLSI, Wayne, PA.

18. van den Bijllaardt W, Schijffelen MJ, Bosboom RW, Cohen Stuart J, Diederen B, Kampinga G, Le TN, Overdevest I, Stals F, Voorn P, Waar K, Mouton JW, Muller AE. 2018. Susceptibility of ESBL Escherichia coli and Klebsiella pneumoniae to fosfomycin in the Netherlands and comparison of several testing methods including Etest, MIC test strip, Vitek2, Phoenix and disc diffusion. J Antimicrob Chemother 73:2380–2387.

19. Camarlinghi G, Parisio EM, Antonelli A, Nardone M, Coppi M, Giani T, Mattei R, Rossolini GM. 2019. Discrepancies in fosfomycin susceptibility testing of KPC-producing Klebsiella pneumoniae with various commercial methods. Diagn Microbiol Infect Dis 93:74–76.

20. Sheppard AE, Stoesser N, Wilson DJ, Sebra R, Kasarskis A, Anson LW, Giess A, Pankhurst LJ, Vaughan A, Grim CJ, Cox HL, Yeh AJ, Sifri CD, Walker AS, Peto TE, Crook DW, Mathers AJ, Group MMMMI. 2016. Nested Russian Doll-like Genetic Mobility Drives Rapid Dissemination of the Carbapenem Resistance Gene blaKPC. Antimicrob Agents Chemother.

21. Clinical Laboratory and Standards Institute. 2018. Performance Standards for Antimicrobial Disk Susceptibility Tests, vol M02. CLSI, 950 West Valley Rd Wayne, PA.

22. Clinical Laboratory and Standards Institute. 2018. Methods for Dilution Antimicrobial Susceptibility Tests for Bacteria That Grow Aerobically, vol M-07, p 91. Clinical Laboratory and Standard Institute, Wayne, PA.

23. Bankevich A, Nurk S, Antipov D, Gurevich AA, Dvorkin M, Kulikov AS, Lesin VM, Nikolenko SI, Pham S, Prjibelski AD, Pyshkin AV, Sirotkin AV, Vyahhi N, Tesler G, Alekseyev MA, Pevzner PA. 2012. SPAdes: a new genome assembly algorithm and its applications to single-cell sequencing. J Comput Biol 19:455–77.

24. Feldgarden M, Brover V. 2019. Using the NCBI AMRFinder tool to determine Anitmicrobial Resistance Genotype-Phenotype correlations within a collection of NARMS Isolates., bioRxiv doi: https://doi.org/10.1101/550707.

25. Eddy SR. 2011. Accelerated Profile HMM Searches. PLoS Comput Biol 7:e1002195.

26. Duez J-M, Mousson C, Siebor E, Pechinot A, Freysz M, Sixt N, Bador J, Neuwirth C. 2011. Fosfomycin and Its Application in the Treatment of Multidrug-Resistant *Enterobacteriaceae Infections*, vol 3, p 123–142. Libertas Academica, Clinical Medicine Reviews in Therapeutics.

27. Zayyad H, Eliakim-Raz N, Leibovici L, Paul M. 2017. Revival of old antibiotics: needs, the state of evidence and expectations. Int J Antimicrob Agents 49:536–541.

28. Testing TECoAS. 2019. Breakpoint tables for interpretation of MICs and zone diameters, version 9.0, 2019., http://www.eucast.org/clinical_breakpoints/.

29. Ito R, Tomich AD, McElheny CL, Mettus RT, Sluis-Cremer N, Doi Y. 2017. Inhibition of Fosfomycin Resistance Protein FosA by Phosphonoformate (Foscarnet) in Multidrug-Resistant Gram-Negative Pathogens. Antimicrob Agents Chemother 61.

30. Guo Q, Tomich AD, McElheny CL, Cooper VS, Stoesser N, Wang M, Sluis-Cremer N, Doi Y. 2016. Glutathione-S-transferase FosA6 of Klebsiella pneumoniae origin conferring fosfomycin resistance in ESBL-producing Escherichia coli. J Antimicrob Chemother 71:2460–5.

31. Rehman MA, Yin X, Persaud-Lachhman MG, Diarra MS. 2017. First Detection of a Fosfomycin Resistance Gene,. Antimicrob Agents Chemother 61.

